# Genomic Analysis of *Salmonella enterica* from cattle, beef and humans in the Greater Tamale Metropolis of Ghana

**DOI:** 10.1101/2024.10.03.616133

**Authors:** Gabriel Temitope Sunmonu, Courage Kosi Setsoafia Saba, Erkison Ewomazino Odih, Opoku Bright, Eric Edem Yao Osei, Alfred Mensah, Saeed Abdallah, Abdul-Razak Alhassan, Stephen Wilson Kpordze, Olabisi C Akinlabi, Anderson O Oaikhena, Beverly Egyir, Iruka N Okeke

**Affiliations:** Department of Pharmaceutical Microbiology, Faculty of Pharmacy, University of Ibadan, Nigeria; Department of Microbiology, Faculty of Biosciences, University for Development Studies, Tamale, Ghana; Department of Biotechnology and Molecular Biology, Faculty of Biosciences, University for Development Studies, Tamale, Ghana; Department of Bacteriology, Noguchi Memorial Institute for Medical Research, University of Ghana, Accra-Ghana

## Abstract

*Salmonella enterica* is a bacterial foodborne pathogen notorious for infecting humans and animals. Proper control of *Salmonella* requires routine surveillance and interventions across the food-production chain. However, due to limited resources the dynamics and transmission of non-typhoidal *Salmonella* serotypes remain poorly understood in several African settings, including within Ghana. Here, we employed bacterial culture and whole genome sequencing (WGS) to investigate the prevalence, virulence and antimicrobial resistance determinants of *Salmonella enterica* isolates from beef, cattle blood and human patient stool in Greater Tamale Metropolis, Ghana. Enrichment and culture of the specimens yielded 62 isolates in total from beef (31), bovine blood (28) and human diarrhoeal specimens (3). We identified at least 15 STs and 18 different *Salmonella* serovars. The most common serovars detected were Poona (n=13), Montevideo (n=10) and Poano (n=7) with S. Montevideo being the most common from cattle blood. Thirty-two isolates belonged to novel sequence types (STs), with ST2609 (n=9) being most common. Four raw beef isolates harboured at least one gene conferring resistance to beta-lactam (*bla*_TEM-1_), chloramphenicol (*catA*), fosfomycin (*fosA7*), quinolone (*qnrD1*) or tetracycline (*tet*(A)). Eight isolates carried at an IncF, IncI and/orCol3M plasmid replicon. This study recovered *Salmonella*, often belonging to previously undocumented STs, at high frequencies from cattle and beef and demonstrated that isolates from human diarrhoeal patients are closely related to bovine isolates. The data highlight the need for broader and sustained surveillance and the urgent need for food safety interventions in Ghana.

## INTRODUCTION

*Salmonella enterica* is a well-known bacterial foodborne pathogen that causes a wide spectrum of infections in both humans and animals(1). The enormous and global impact of *Salmonella* on public health and food safety stems from their aetiologic role in gastroenteritis, bacteraemia, and systemic infections (2, 3). Many cases of non-typhoidal *Salmonella* in humans are of zoonotic origin and attributable to Subspecies 1 of *Salmonella enterica* (4).

The incidence of *Salmonella* from meat and other animal products can vary depending factors such as adherence to food safety regulations, animal infection incidence, handling, cooking methods, and epidemiological surveillance systems in place (5, 6). *Salmonella* contamination of animal products remains a significant concern for public health worldwide. In the United States, European Union member states, and many other parts of the world, regulatory agencies monitor and track foodborne illnesses, including those caused by *Salmonella*, through surveillance programs (7). These programs help to estimate the incidence of *Salmonella* infections associated with beef consumption.

Despite the high incidence of both gastrointestinal and invasive non-typhoidal *Salmonella* infections (8, 9), *Salmonella enterica* remains understudied across Africa. Previously reported *Salmonella* serovars in Ghana include Kaapstad (ST4605 from beef and mutton), Hato (ST5308 from chicken; ST3899 from guinea fowl), Lagos (ST2469 from beef and goat) and Infantis (ST603 from goat) (10). Additionally, *Salmonella* serovars identified in chicken eggs in Ghana include S. enteritidis (ST11), S. Hader, and S. Chester. These strains carry virulence genes, such as those encoding fimbrial adherence and secretion systems (*inv, spa, ssa and sse*), and resistance determinants against fluoroquinolones (*gyrA* (D87N), *qnrB81*), aminoglycosides (*aadA1, aph(3”)-Ib aph(6)-Id*), tetracycline (tet(A)), phenicols (*catA1*) and trimethoprim (*dfrA14 and dfrA1*) (11). While some studies (10-12) have investigated the role of meat and other animal products in the spread of Salmonellosis in Ghana, no study has investigated its prevalence in slaughter cattle. Understanding the transmission dynamics and genetic diversity of *Salmonella enterica* across different reservoirs is crucial for devising effective control strategies and mitigating its associated risks. In this study, we determined the prevalence of *Salmonella enterica* in the beef value chain and among humans presenting with diarrhoea in hospitals in the Greater Tamale Metropolis of Ghana.

## METHODS

### Experimental Design and Sample Collection

Beef samples were collected randomly from vendors within the Greater Tamale Metropolis. Cattle blood samples were also randomly collected at the Greater Tamale Abattoir from. These cattle are brought from a range of locations In Ghana, as well as from other countries, including Burkina Faso, Mali, and Niger, to the Greater Tamale Abattoir where they are slaughtered and processed for sale. 30-35 ml of cattle blood samples were collected aseptically into labelled sterile 50 ml tubes (Falcon®). Faecal samples were collected from patients with diarrhoea or gastroenteritis cases attending the Tamale Central Hospital, Tamale West Hospital and Seventh Day Adventist Hospital using sterile stool containers. All samples (beef, cattle blood and human faecal sample) were transported in ice chests with ice within 2 hrs of collection to the Spanish Laboratory Complex of the University for Development Studies, Tamale, Ghana for analyses.

### Salmonella isolation and identification

*Salmonella enterica* were isolated on Xylose Deoxycholate (XLD) agar after pre-enrichment on peptone and enrichment on Modified Semi-Solid Rappaport-Vassiliadis (MSRV) agar. Bacterial colonies having morphological characteristics similar to *Salmonella* were sub-cultured to obtain pure colonies and Gram stained. PCR targeting the invA gene as described by Rahn et al. (1992) (13), using PCR oligonucleotides using PCR oligonucleotides invA139f GTGAAATTATCGCCACGTTCGGGCAA and *invA141r* TCATCGCACCGTCAAGGAACC. Previously described PCR protocol was used (14). PCR products were separated on 1.5% (w/v) agarose gels, stained with gel red (Biotium) and visualized using a transilluminator (UVP GelMax Imager).

Further biochemical profiling of isolates and biotyping, was conducted using the Gram-negative (GN) test kit (Ref: 21341) on VITEK 2 systems (version 2.0, Marcy-l*’*Etoile, France, Biomérieux) according to manufacturer*’*s instructions. Strains that identified as *Salmonella* biochemically, were whole genome sequenced.

### DNA Extraction, Library Preparation and Whole Genome Sequencing

Genomic DNA of the isolates were extracted using Wizard DNA extraction kit (Promega; Wisconsin, USA) in accordance with manufacturer*’*s protocols. Using a dsDNA Broad Range quantification assay, the extracted DNA was quantified on a Qubit fluorometer (Invitrogen; California, USA). DNA libraries were prepared using NEBNext Ultra II FS DNA library kit for Illumina with 96-unique indexes (New England Biolabs, Massachusetts, USA; Cat. No: E6609L). DNA libraries was quantified using dsDNA High Sensitivity quantification assay on a Qubit fluorometer (Invitrogen; California, USA). The average fragment length of the DNA libraries was determined using 2100 Bioanalyzer (Agilent). Libraries were sequenced on an Illumina MiSeq (Illumina, California, USA). The raw sequence reads were de novo assembled using SPAdes v3.15.3 (15) as implemented in the GHRU assembly pipeline (https://gitlab.com/cgps/ghru/pipelines/dsl2/pipelines/assembly).

### Sequence Typing of Salmonella Genomes

Sequence reads were deposited in the *Salmonella* database for *Salmonella* on EnteroBase (16) to determine the multi-locus sequence types (MLST) and core-genome MLSTs. The *Salmonella* genome assemblies were analysed using the *Salmonella* In-Silico Typing Resource (SISTR) for the prediction of serovars and serogroups (17) (https://github.com/phac-nml/sistr_cmd).

### Identification of Plasmids, Antimicrobial resistance (AMR) and Virulence genes

Plasmid replicons present in the assembled genomes were detected using PlasmidFinder (18). Antimicrobial resistance genes carried by the isolates and the drug classes to which they possibly conferred resistance were predicted using AMRFinderPlus v3.10.24 (19) and its associated database (version 2022-04-04.1). We also detected the virulence genes present in the isolates using ARIBA (20) with the virulence factor database (VFDB).

### Single Nucleotide Polymorphism (SNP) Calling and Phylogenetic analysis

For phylogenetic analysis, reference sequences for the *S. enterica* genomes were objectively selected from the National Center for Biotechnology Information Reference Sequence (RefSeq) database (https://www.ncbi.nlm.nih.gov/refseq/) using BactinspectorMax v0.1.3 (https://gitlab.com/antunderwood/bactinspector). The selected reference was the *S. enterica* subsp. enterica serovar Poona strain (assembly accession: GCF_900478385.1). The sequence reads for each species were then mapped to the chromosome of the reference using BWA (v0.7.17) (21) and variants were called and filtered using BCFtools v1.9 (22) as implemented in the GHRU single nucleotide polymorphism (SNP) phylogeny pipeline (https://gitlab.com/cgps/ghru/pipelines/snp_phylogeny). Variant positions were concatenated into a pseudoalignment and used to generate a maximum likelihood tree using IQ-TREE v1.6.8 (23). SNP distances between the genome pairs were calculated using snp-dists v.0.8.2 (https://github.com/tseemann/snp-dists).

### Ethical considerations

Ethical approval to sequence the isolates in this study for research and surveillance was obtained from the Noguchi Memorial Institute for Medical Research NIMIMP-052/21/22

### Data availability

The sequence reads were deposited in the European Nucleotide Archive with the study accession PRJEB58695 (Supplementary Table 1).

**Table 1:** Overview of all 62 isolates and associated ST, serotype, resistance genes and plasmid replicons.

| ID | ST | serotype | Resistance Genes | Plasmid Replicons |
| --- | --- | --- | --- | --- |
| GH-CKS-105B_S5 | 306 | Bredeney | *qnrD1* | Col3M, IncI1-I(Gamma) |
| GH-CKS-106B_S6 | 308 | Poona | - | - |
| GH-CKS-124B_S12 | 308 | Poona | *bla*<sub>TEM-1</sub>,<br>*tet(A)* | IncFIB(AP001918),<br>IncFII, IncI2(Delta) |
| GH-CKS-74B_S4 | 321 | Muenster | - | IncFII(S) |
| GH-CKS-18B_S24 | 411 | Chester | - | - |
| GH-CKS-BC-92_S11 | 411 | Chester | - | - |
| GH-CKS-BC98_S2 | 411 | Chester | - | - |
| GH-CSK-37B_S1 | 411 | Chester | - | - |
| GH-CKS-BC-68_S23 | 516 | Give | - | - |
| GH-CKS-BC-72_S21 | 516 | Give | - | - |
| GH-CKS-BC-73_S16 | 516 | Give | - | - |
| GH-CKS-BC71_S36 | 516 | Give | - | - |
| GH-CKS-92B_S24 | 2152 | Gaminara | - | - |
| GH-CKS-CH-13_S9 | 2152 | Gaminara | - | - |
| GH-CKS-BC61_S28 | 2152 | Gaminara | - | - |
| GH-CSK-87B_S29 | 2176 | Labadi | - | - |
| GH-CKS-23B_S10 | 2609 | Poona | - | - |
| GH-CKS-47B_S8 | 2609 | Poona | - | IncFII(SARC14) |
| GH-CKS-BC-69_S19 | 2609 | Poona | - | IncFII(S) |
| GH-CKS-BC-70_S20 | 2609 | Poona | - | - |
| GH-CSK-65B_S27 | 2609 | Poona | - | IncFII(SARC14) |
| GH-CSK-BC62_S3 | 2609 | Poona | - | - |
| GH-CSK-BC65_S2 | 2609 | Poona | - | - |
| GH-CSK-BC67_S4 | 2609 | Poona | - | - |
| GH-CSK-BC78_S5 | 2609 | Poona | - | IncFII(S) |
| GH-CKS-BC-60_S22 | 2609* | Poona | - | - |
| GH-CKS-103B_S4 | 6829 | Pisa | - | - |
| GH-CKS-2B_S2 | 7643 | Scarborough | *fosA7* | - |
| GH-CKS-11B_S17 | 8515 | Odozi | - | - |
| GH-CSK-100B_S25 | 8515 | Odozi | - | - |
| GH-CKS-28B_S27 | 8515* | Odozi | - | - |
| GH-CKS-21B_S16 | 8631 | Vinohrady | - | - |
| GH-CKS-114B_S43 | 10435* | Chester | - | - |
| GH-CKS-20B_S41 | 10437* | Poano | - | - |
| GH-CKS-27B_S9 | 10438* | Yoruba | - | - |
| GH-CKS-BC-79_S18 | 10439* | Poano | - | - |
| GH-CKS-BC-80_S23 | 10439* | Poano | - | - |
| GH-CKS-BC-90_S11 | 10439* | Poano | - | - |
| GH-CKS-BC64_S6 | 10439* | Poano | - | - |
| GH-CSK-BC88_S7 | 10439* | Poano | - | - |
| GH-CKS-115B_S7 | 10436* | Llandoff | - | - |
| GH-CKS-13B_S3 | 10436* | Llandoff | - | - |
| GH-CKS-3B_S15 | 10463* | I 1,3,19:z:- | - | - |
| GH-CKS-4B_S14 | 10463* | I 1,3,19:z:- | - | - |
| GH-CKS-67B_S4 | 10462* | I 28:g,t:- | - | IncFII(S) |
| GH-CKS-78B_S5 | 10440* | Muenster | - | - |
| GH-CKS-83B_S6 | 10436* | Llandoff | - | - |
| GH-CKS-84B_S10 | 10462* | I 28:g,t:- | - | - |
| GH-CKS-88B_S13 | 10461* | I 4:y:- | *catA* | - |
| GH-CKS-BC-100_S8 | 10441* | Montevideo | - | - |
| GH-CKS-BC-77_S17 | 10441* | Montevideo | - | - |
| GH-CKS-BC-85_S15 | 10441* | Montevideo | - | - |
| GH-CKS-BC-93_S12 | 10441* | Montevideo | - | - |
| GH-CKS-BC74_S37 | 10441* | Montevideo | - | - |
| GH-CKS-BC79B_S12 | 10441* | Montevideo | - | - |
| GH-CKS-BC81_S38 | 10441* | Montevideo | - | - |
| GH-CKS-BC84_S1 | 10441* | Montevideo | - | - |
| GH-CKS-BC87_S8 | 10439* | Poano | - | - |
| GH-CKS-CH-12_S7 | 10441* | Montevideo | - | - |
| GH-CKS-CH-24_S10 | 10441* | Montevideo | - | - |
| GH-CSK-71B_S28 | 10440* | Muenster | - | - |
| GH-CKS-BC56_S33 | 10442* | Poona | - | - |

Key: \* = Novel STs. Supplementary Table 2 has additional details on the isolates

## RESULTS

### Prevalence of *Salmonella* in the studied specimens

We collected and processed a total of 124 beef samples, 150 cattle blood samples, and 80 human stool samples. Culture and initial tests on MSRV and XLD yielded presumptive *Salmonella* from 37/124 (29.8%) raw beef samples, of which 31 (25% of the 124 specimens) were eventually confirmed as *S. enterica* based on whole genome sequencing (WGS). For the cattle blood samples, 37/150 (24.7%) were positive for presumptive *Salmonella*, and 28/150 (18.7%) were confirmed positive. Among the human faecal samples, recorded 3/80 (3.8%) were positive for presumptive *Salmonella*, all of which were confirmed to be *S. enterica* based WGS. In all, the genomes of sixty-two *Salmonella* isolates were sequenced in this study; 31 were from raw beef, 28 from blood of slaughtered cattle, and three from human stool.

### Salmonella Serotypes, Sequence Types (STs), and Phylogeny

The most common serotypes detected in this study were Poona (n=13, 21%), Montevideo (n=10, 16.1%), Poano (n=7, 11.3%), Chester (n=5, 8.1%) and Give (n=4, 6.5%). Other represented serotypes include; Gaminara (n=3), Llandoff (n=3), Muenster (n=3), Odozi (n=3), Bredeney (n=1), Labadi (n=1), Pisa (n=1), Scarborough (n=1), Vinohrady (n=1), Yoruba (n=1), I 1,3,19:z:-(n=2), I 28:g,t:-(n=2) and I 4:y:-(n=1) (Table 1).

The most common STs were ST2609 and ST411 which were represented by ten (14.5%) and four (6.5%) isolates respectively (Figure 1). As many as 32 (51.6%), including all 10 Montevideo (ST10441) and seven isolates of the rare Poano serotype (ST10439 (n=6), ST10437 (n=1)). The 13 S. Poona isolates belonged to ST2609 (n=10), ST308 (n=2), novel ST10442 (1). Four of the Chester serotype strains belong to ST411 while one is a single locus variant of the sequence type (ST411*). The four Give serotype belong to ST516 (Figure 1, Table 1).

**Figure 1.**
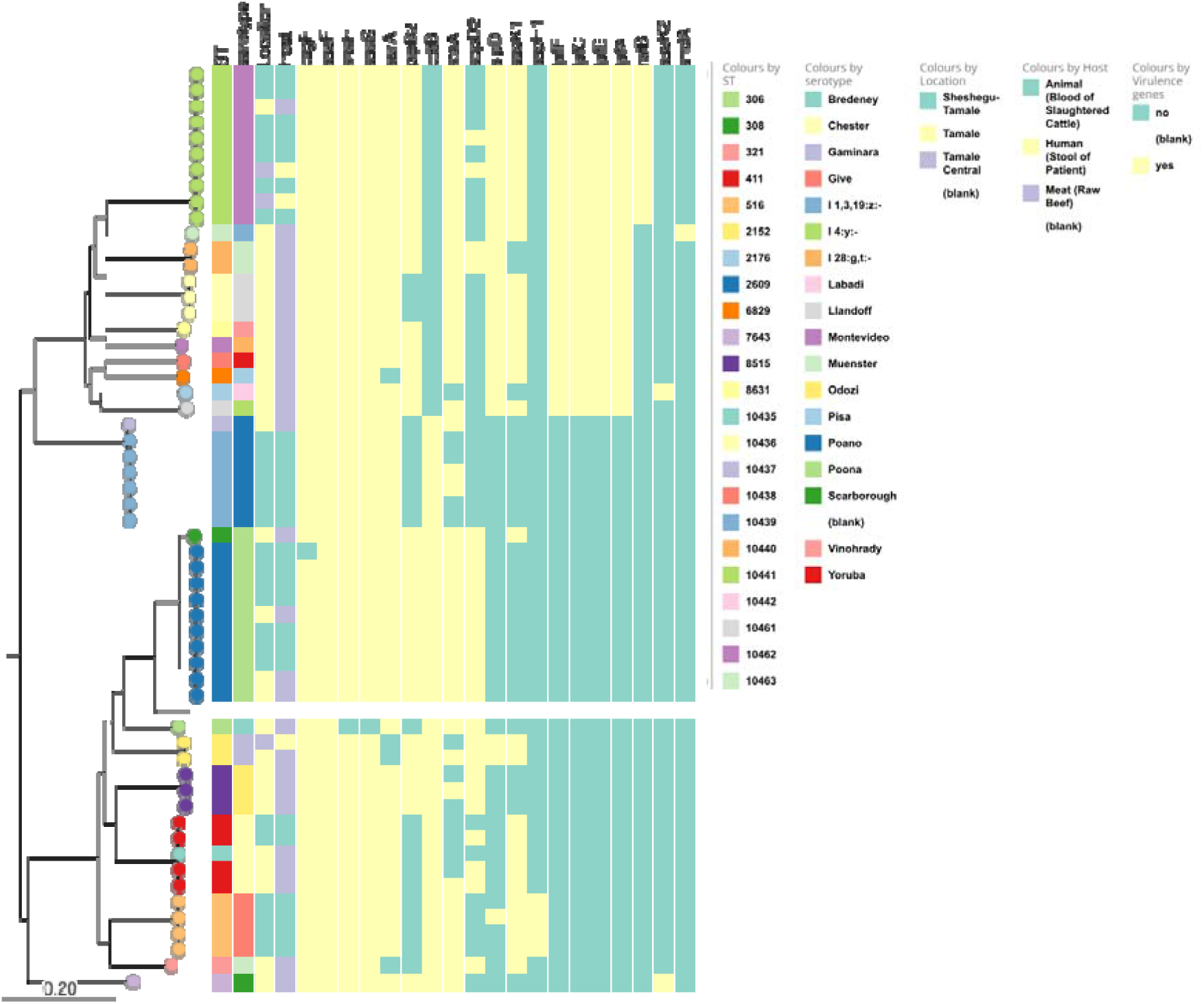
Phylogeny of *Salmonella* isolates. A mid-point rooted maximum likelihood phylogenetic tree based on SNP alignment and created using IQ-Tree annotated with ST, serotypes, location of isolation, host of isolates and virulence genes not shared by all isolated in the assembled genomes.

The most diversity in serotypes and sequence types seen in this study was seen among raw beef and slaughtered cattle blood isolates. Uncommonly recovered ST306 (n=1), ST308 (n=2), ST321 (n=1), ST2176 (n=1), ST7643 (n=1), ST8515 (n=2) and ST8631 (n=1) were found only in raw beef isolates and ST516 (n=4) was found only in blood of slaughtered cattle. Two of the human stool isolates belong to Montevideo serotype and a previously unreported sequence type (ST10441) while the third belongs to ST2152 and the Gaminara serotype (Figure 1, Table 1). Both of these serotypes and STs were also recovered from bovine sources in the study.

A SNP-based phylogeny of 57 *Salmonella* isolates from this study is presented in Figure 1. Most of the isolates belong to the most prominent *Salmonella* enterica clades A (including the Pisa and Yoruba isolates) and B (including the Give and Chester isolates). Isolates belonging to the same serovar and sequence type cluster together on the phylogenetic tree except for the three S. Muenster strains. Two Novel ST (ST10440) S. Muenster strains cluster together in Clade A while the third belonging to ST321 was located in Clade B on the tree, consistent with the fact that some S. enterica serovars (including Muenster and Montevideo) are comprised of strains from distant clades). The two human and six of the animal *S*. Montevideo isolates had no SNP difference. The remaining two animal *S*. Montevideo isolates are two SNP differences apart and one SNP difference from the rest. One *S*. Gaminara isolate from human stool appears to be closely related to another from raw beef with these two strains showing only 53 SNP differences (Supplementary Table 3).

### Virulence Factors, Plasmid Replicons and ARG Profiles of Salmonella

All the isolates carry multiple genes that have been shown to be important for *Salmonella* virulence. The curli (csg) genes were present in all *Salmonella* isolates as well as the fimbrial operon *bcf*, and *fim*) genes. All the isolates had at least two ste fimbrial genes, *steB* and *steC*. Isolates, belonging to all serovars except *S*. Bredeney, Chester, Gaminara Give, Odozi, Poano, Poona and Scarborough, 38.7% (24/62), carried long polar fimbriae (*lpf*) genes. Only the *S*. Montevideo strains carried the virulence factor *ratB*, associated with long-term intestinal persistence and the I 1,3,19:z:-strain carry *shdA*, which encodes an autotransporter. The *inv, org, prg, sif, spa, ssa, ssc, sse, ssp* and *sop* type III secretion system effector genes were detected in all the isolates while *avr* was present in 91.9% (57/62) of the isolates. Thirty-eight (61.3%) of the isolates also carry the cytholethal distending toxin gene, *cdtB*. Genes involved in host cell manipulation and invasion such as: *slrP, sptP, sicP, sicA, pipB, mig-14, misL, sinH* and *sseL* were present in all isolates while *pipB2* was present in 66.1% (41/62) of isolates (Figure 1, Supplementary Table 4). The virulence genes *gogB* and *grvA* were absent in all strains.

Most of the *Salmonella enterica* isolates (58/62) harbour no AMR genes. However, four isolates from raw beef carry at least one of the gene conferring resistance to beta-lactam (*bla*_TEM-1_), chloramphenicol (*catA*), fosfomycin (*fosA7*), quinolone (*qnrD1*) and tetracycline (*tet(A))* (Table 1).

The plasmid replicons identified in this study were IncFII(S) (n=4), IncFII(SARC14) (n=2), IncFII (n=1), IncFIB(AP001918) (n=1), IncI2(Delta) (n=1), Col3M (n=1) and IncI1-I(Gamma) (n=1). The plasmid replicons were spread across eight isolates (6 raw beef and 2 blood of slaughtered cattle isolates). Four of the ST2609 serovar Poona isolates harbour one of either IncFII(S) or IncFII(SARC14) while one raw beef isolate (GH-CKS-124B_S12) harbour three plasmid replicons (IncFII(SARC14), IncFII and IncI2(Delta)) (Table 1).

## DISCUSSION

*Salmonella* is the aetiologic agent of foodborne illnesses worldwide, including in Ghana (11, 12). Meat is often implicated in Salmonellosis (24). *Salmonella* can be transmitted from infected cows and, more commonly, butchery cross-contamination can occur through a range of pathways, including via contaminated surfaces, improper raw meat handling, poor employee hygiene, equipment contamination, and incorrect storage practices (25). In this study, *Salmonella* were recovered from over 20% of the slaughter cattle blood and meat sampled. We sequenced and analysed the genomes of 59 bovine isolates and three isolates recovered from human stool samples.

A few *S*. enterica serovars that are commonly reported globally were detected in this study, including Montevideo, Muenster and Poona but most the serotypes detected were uncommon in the literature overall (eg *S*. Poano and Odozi) or commonly reported in African, rather than in other settings (eg *S*. Give and Yoruba) (26-29). A number of STs belonging to these and more commonly reported serovars were novel. Most of the *Salmonella* serovars identified in this study have not yet been reported in Ghana except for AMR *S*. Poona isolated from poultry meat (30) and humans (31). *S*. Gaminara has been associated with citrus and other fruits (32-35) and water (36, 37) whereas in this study, one human stool and raw beef isolate was identified as *S*. Gaminara. Isolation of S. Montevideo has also been reported in buffalo meat in Egypt (38), in both cattle and human in the US (39), in non-diarrhoeic dogs in Grenada, West Indies (40) and in dairy farms in Uruguay (41). Similarly, in this study *S*. Montevideo was present in human stool, raw beef and blood of slaughter cattle.

The presence of strains belonging at least 15 STs and 18 different *Salmonella* serovars in cattle blood, raw beef and human stool from this study, demonstrates the diversity of Salmonella lineages circulating in the beef foodchain and the need for preventive actions. These should include improved hygiene and sanitation practices, strengthened surveillance and monitoring systems, stricter food safety standards and regulations, vaccination and treatment programs for cattle, and public health awareness campaigns. Such actions would help reduce the transmission of foodborne pathogens and lower the risk of related illnesses. Most of these serovars and STs have not been previously documented in Ghana, highlighting the need for more and more precise surveillance, as can be accomplished through whole genome sequencing.

The IncFII(S) plasmid replicon was detected more commonly than any other in *Salmonella* in this study. Plasmids with this replicon have been reported to carry genes that confer resistance to beta-lactams (42-45), tetracyclines (42, 45) and aminoglycosides (42) as well as virulence (42, 43) in *Salmonella*. Other plasmid replicons detected in this study such as: IncFII(SARC14) (46), IncFII (45, 47), IncFIB(AP001918) (45), IncI2(Delta), Col3M (48) and IncI1-I(Gamma) (49) have been reported to carry AMR genes in *Salmonella*. However, in this study none of the isolates carrying IncFII(S) and IncFII(SARC14) replicons harbour any resistance genes. This may reflect a lack of selective pressure in the beef production sector. However the presence of replicons associated with resistance means that platforms are in place for rapid evolution to resistance. The *S*. Bredeney ST306 isolate harbouring the plasmid-mediated quinolone resistance gene *qnrD1* carries Col3M and IncI1-I(Gamma) plasmid replicons and one of the S. Poona ST308 isolates (GH-CKS-124B_S12) carrying IncFII, IncFIB(AP001918) and IncI2(Delta) also harbours tetracycline (*tet(A))* and beta-lactamase (*bla*_TEM-1_) resistance genes. The IncI2 replicon is commonly reported from E. coli, where it carries multiple AMR genes (50), but rarely present in *Salmonella* and when it is detected, does not usually harbour any resistance determinants (41, 51).

The most concerning finding in this study is that *Salmonella* was recovered from three of 80 stool specimens and that all three isolates were closely related to bovine isolates. All three strains belonged to serovars recovered from beef – Gaminara and Montevideo – and S. Montevideo isolates were also the most common serovar of slaughter cattle blood isolates. Human and bovine isolates in all instances were highly similar., The data in this study suggest interchange of *Salmonella* between humans and livestock/meat.

## CONCLUSION

This study has identified a range of *Salmonella* serovars in slaughter cattle and beef in Ghana. The serovariety types and virulence gene repertoires of the recovered strains suggest pathogenic potential and recovery of strains belonging to two lineages identified from humans visiting health centres in the region further gives cause for concern. While antimicrobial resistance was not common in this study, resistance genes and mobile elements capable of carrying resistance genes were identified in many of the isolates, emphasizing the need to avoid transmission of *Salmonella* across One Health boundaries in Ghana. To effectively combat the spread of *Salmonella*, it is essential to foster collaboration among various stakeholders in the One Health framework, including public health officials, veterinarians, food safety experts, and environmental scientists, thus creating a comprehensive and integrated approach to AMR surveillance and control.

## Supplementary Materials

Supplementary Table 1: Accession IDs, Supplementary Table 2: Isolates*’* STs and allelic profile Supplementary Table 3: SNP distance matrix, Supplementary Table 4: Isolates*’* metadata including plasmid replicon, AMR and virulence profile.

## Acknowledgements

We thank Ayorinde Afolayan, Faith I. Oni, and Abeeb Adeniyi for technical assistance and Jola-Ade Ajiboye, Kesiena Akpede and Pernille Nilsson for logistic support.Reagents and media for the initial isolation and identification were supported by Bruno Gonzalez-Zorn of the Antimicrobial Resistance Unit of the Universidad Complutense de Madrid, Spain. Whole genome sequencing and analysis for this project was supported through SEQAFRICA. The SEQAFRICA project is funded by the Department of Health and Social Care*’*s Fleming Fund using UK aid. INO is a Calestous Juma Fellow supported by the Bill and Melinda Gates Foundation. The views expressed in this publication are those of the authors and not necessarily those of the UK Department of Health and Social Care, its Management Agent, Mott MacDonald or any of the other funders.

